# Lifestyle factors in the biomedical literature: An ontology and comprehensive resources for named entity recognition

**DOI:** 10.1101/2024.06.13.598816

**Authors:** Esmaeil Nourani, Mikaela Koutrouli, Yijia Xie, Danai Vagiaki, Sampo Pyysalo, Katerina Nastou, Søren Brunak, Lars Juhl Jensen

**Affiliations:** Novo Nordisk Foundation Center for Protein Research, Faculty of Health and Medical Sciences, University of Copenhagen; Faculty of Information Technology and Computer Engineering, Azarbaijan Shahid Madani University, Tabriz, Iran; TurkuNLP group, Department of Computing, Faculty of Technology, University of Turku, Turku, Finland

**Author notes:** Zhejiang Ponshine Information Technology Co., Ltd., Hangzhou 311100, China (Yijia Xie). a. Genome Biology Unit, European Molecular Biology Laboratory (EMBL), Heidelberg, Germany, b. Collaboration for joint PhD degree between EMBL and Heidelberg University, Faculty of Biosciences, Heidelberg, Germany, c. Division of Computational Genomics and Systems Genetics, German Cancer Research Center (DKFZ), Heidelberg, Germany.

## Abstract

**Motivation:** Despite lifestyle factors (LSFs) being increasingly acknowledged in shaping individual health trajectories, particularly in chronic diseases, they have still not been systematically described in the biomedical literature. This is in part because no named entity recognition (NER) system exists, which can comprehensively detect all types of LSFs in text. The task is challenging due to their inherent diversity, lack of a comprehensive LSF classification for dictionary-based NER, and lack of a corpus for deep learning-based NER.

**Results:** We present a novel Lifestyle Factor Ontology (LSFO), which we used to develop a dictionary-based system for recognition and normalization of LSFs. Additionally, we introduce a manually annotated corpus for LSFs (LSF200) suitable for training and evaluation of NER systems, and use it to train a transformer-based system. Evaluating the performance of both NER systems on the corpus revealed an F-score of 64% for the dictionary-based system and 76% for the transformer-based system. Largescale application of these systems on PubMed abstracts and PMC Open Access articles identified over 300 million mentions of LSF in the biomedical literature.

**Availability:** LSFO, the annotated LSF200 corpus, and the detected LSFs in PubMed and PMC-OA articles using both NER systems, are available under open licenses via the following GitHub repository: https://github.com/EsmaeilNourani/LSFO-expansion. This repository contains links to two associated GitHub repositories and a Zenodo project related to the study. LSFO is also available at BioPortal: https://bioportal.bioontology.org/ontologies/LSFO.

**Supplementary information:** Supplementary data are available at *Bioinformatics* online.

## 1 Introduction

Lifestyle is a complex and multifaceted concept that involves nutrition, behaviors, habits, and activities that individuals engage in, which can impact their health. These lifestyle factors (LSFs) are widely recognized as important in shaping individual health trajectories and influencing the onset and progression of diseases (WHO, 2023). Numerous studies illustrate the association between an unhealthy lifestyle and an elevated risk of chronic disease as well as the potential of reducing the risk by adopting a healthy lifestyle (Subramanian *et al*., 2020; Nyberg *et al*., 2020; Tobias *et al*., 2023). LSFs can thus play a pivotal role together with genetic factors in precision medicine (Yurkovich *et al*., 2023; Gabbert *et al*., 2023; Gray *et al*., 2020; Jeon *et al*., 2018), and a crucial first step is to structure the existing information on LSFs.

Specific types of LSFs such as environments, exposure to risk factors, smoking, and food are covered by existing resources like the Environment Ontology (Buttigieg *et al*., 2013), the Exposome Explorer (Neveu *et al*., 2020), the Cigarette Smoke Exposure Ontology (Younesi *et al*., 2014), and FoodOn (Dooley *et al*., 2018). However, these fall short in comprehensively covering other crucial disease-associated LSFs. Since, there is no clear definition in the literature of what constitutes an LSF, we define LSFs to be all non-genetic health determinants associated with diseases. This includes categories like *physical and leisure time activities, socioeconomic factors, personal care products and cosmetic procedures, sleep, mental health practices and substance use*, on top of the categories mentioned above. The fact that there is no comprehensive ontology covering all these categories severely hampers formalization of information on LSFs, including the development of text-mining solutions to help extract it from literature.

The first critical step to alleviate this issue is to develop a named entity recognition (NER) method to identify LSF mentions within scientific publications. Many such systems exist for other biomedical entities, reviewed in (Song *et al*., 2021; Huang *et al*., 2020; Perera *et al*., 2020). Developing an NER method for LSFs requires either a comprehensive LSF dictionary that can be matched against text to find them, or a text corpus with manually annotated LSFs that can be used to train deep learning-based methods. In this paper, we address the task of recognizing LSFs within biomedical text by presenting four major contributions. Firstly, we introduce a novel LSF ontology (LSFO), featuring a multilevel hierarchical structure. It includes the main LSF categories at the top level and extends to specific subcategories and low-level concepts. Secondly, we introduce the first annotated corpus for LSFs, LSF200, a valuable resource for the BioNLP community as it can be used for training and evaluating NER systems. Thirdly, we introduce a dictionary-based NER system enabling for the first time the detection of a diverse set of lifestyles in the literature. This NER system leverages the LSFO for dictionary creation, thus enabling both recognition and normalization for the matches to LSFO concepts. Lastly, we introduce a transformer-based NER system (Vaswani *et al*., 2017) for LSF detection, leveraging LSF200 for training and evaluation.

## 2 Materials and Methods

Currently, there is a lack of resources to capture the vast diversity of LSFs, under a single umbrella. Below, we have made an effort to generate a comprehensive categorization of the different aspects of LSFs accompanied by a small description of concepts that fit in each category. The creation of this categorization is the first crucial step that allowed us to annotate a text corpus and create a dictionary of LSFs, for the purposes of deep learning-based NER and dictionary-based NER, respectively. In the next sections we provide details in the methodology used to achieve our goals.

### 2.1 The LSF categorization

Within the context of lifestyle, we have identified nine categories that can collectively describe all LSFs, namely:

1. **Nutrition**, a category that covers different branches such as dietary habits, food groups, food processing and preparation, macronutrients, and micronutrients, among others.
2. **Socioeconomic factors**, which includes social and economic conditions such as income, wealth, education, and socioeconomic status.
3. **Environmental exposures**, includes exposure to various environmental factors, such as air pollution, water quality, and workplace hazards, that can impact an individual’s health.
4. **Substance use**, covers concepts such as smoking, as well as illicit drug use.
5. **Physical activities**, includes regular exercise and physically demanding activities like leisure time, occupational, and household physical activities.
6. **Non-physical leisure time activities**, describes any activity an individual might engage in during their free time that is not a physical activity.
7. **Personal care products and cosmetic procedures**, covers activities related to hygiene, use of cosmetic and cleaning products as well as invasive procedures that people undergo to improve their appearance, such as cosmetic surgery.
8. **Sleep**, covers sleep quality, stages, and habits.
9. **Mental health practices**, includes the behaviors and habits related to maintaining good mental health and emotional well-being such as meditation, and psychotherapy.

### 2.2 Annotation of the LSF200 text corpus

To create an LSF text corpus, we selected three of the most relevant journals within each of the nine LSF categories introduced above. We selected 200 abstracts for our corpus, evenly distributed across all categories. We aimed for a balanced selection of documents among LSF categories, considering the chosen journals’ scope and focus. Selected top three journals per category are available in Supplementary Section 1. To ensure high diversity, we opted to annotate abstracts instead of full-text documents, as this would allow us to annotate a larger number of documents and thus obtain a wider selection of different entities with the same curation effort. The annotation process started by creating an initial set of annotation guidelines, which we improved through two rounds of refinement. For each of the three versions of the guidelines, two annotators annotated a new set of 15 abstracts, based on which we calculated the inter-annotator agreement. A meeting was held after each round to discuss disagreements, update the guidelines, and clarify any ambiguities or gaps in the rules that caused the disagreements between the annotators. We subsequently annotated the entire LSF200 according to the final guidelines (https://esmaeilnourani.github.io/lifestylefactors-annotation-docs/entities), with each annotator working on a different set of abstracts. We used the BRAT rapid annotation tool for document annotation (Stenetorp et al., 2012)

### 2.3 Dictionary construction

One common approach to NER is to develop a dictionary-based system (Cook and Jensen, 2019), leveraging existing databases or ontologies to create the dictionary. For instance, a disease NER system can be established by constructing a dictionary derived from existing resources like the Disease Ontology (Baron *et al*., 2024). While there are existing ontologies that cover some aspects of LSFs, there is no comprehensive resource that encompasses all aspects; thus, we manually constructed an initial dictionary inspired by existing biomedical literature, ontologies, and questionnaires. The aim was to come up with a wide selection of names belonging to each of the nine categories of LSFs described above, which would serve as a good starting point for semi-automatic expansion. The guidelines for the creation of the manual version of the dictionary are provided in Supplementary Section 2.

Then, we wanted to introduce names from a resource that would contain a diverse set of candidates. As 45% of the names in the initial dictionary exist as Wikipedia pages, we decided to use Wikipedia page titles as a source of potentially missing LSFs. To predict which page titles are good LSF candidates, we used a transformer-based approach that scores names based on their contexts in biomedical literature (Nastou *et al*., 2023), and manually assessed all high-scoring candidates before adding them to the dictionary. For more details on the training, evaluation and prediction process please refer to Supplementary Section 3.

Afterwards, we incorporated names from resources such as WordNet (Brown, 2005), Wikidata, DBpedia (Lehmann *et al*., 2015), ConceptNet (Speer *et al*., 2018), and 1085 existing ontologies registered in BioPortal (Whetzel *et al*., 2011). LSF-related candidates were extracted from these resources, and scored based on both semantic similarity to existing names and textual context using BERTopic (Grootendorst, 2022). The details of this process are provided in Supplementary Section 4.

Figure 1 displays the details of different stages of LSF dictionary creation and expansion, along with methods used and the resources considered in each stage as LSF candidates. Apart from the initial creation of the dictionary, which was entirely performed manually, the subsequent two automatic expansion steps also included manual validation and filtering of candidates, before their addition to the dictionary.

**Fig. 1.**
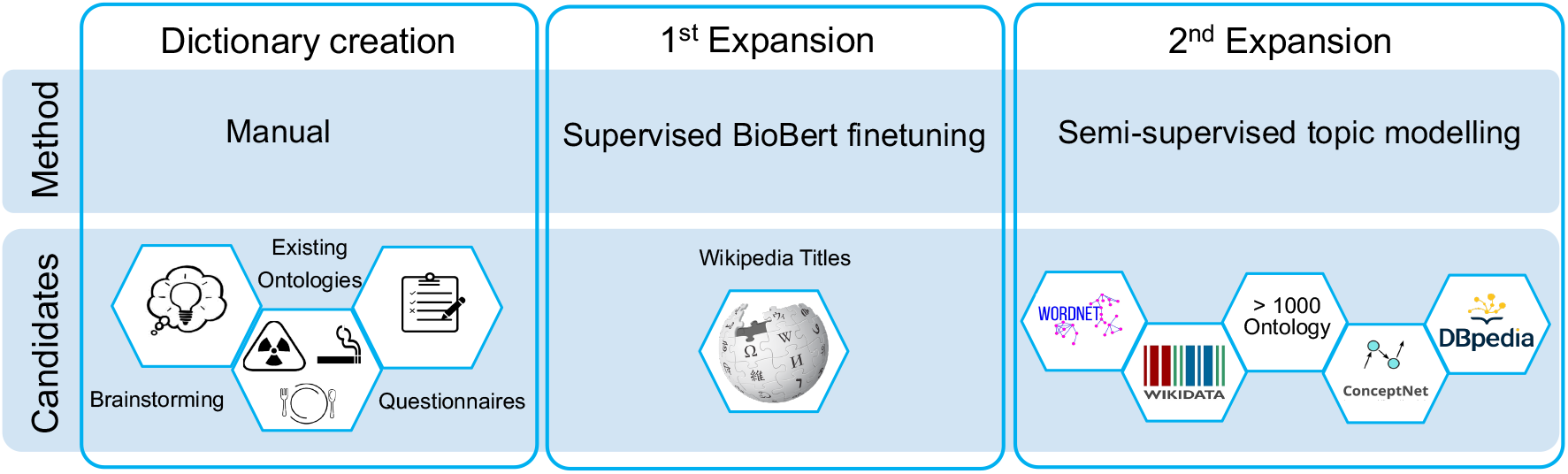
LSF Dictionary creation and expansion stage

### 2.4 Creation of an LSF ontology and referencing to external resources

In this study, we defined nine distinct categories, encompassing diverse LSF aspects, to establish a unified LSF classification across the various categories. For practical uses of the dictionary, such as concept indexing, it is important to know which names are synonyms for the same LSF and how the LSFs relate to each other. We thus created an LSF ontology (LSFO) in which each LSF is a concept with a unique identifier associated with relevant synonyms, and the concept can be traced back to the root of the LSFO through *is_a* relationships. To cover fine-grained concepts related to food groups, we imported the entire Food Groups branch of FoodOn, preserving its hierarchy. We also imported concepts from other ontologies; however, this was done in a selective manner involving manual curation and did not necessarily preserve the hierarchy of the source ontology. Details on the construction of LSFO and the process and tools used for conflict resolution are provided in Supplementary Section 5.

We utilized the BioPortal annotator to match the existing LSFs in our LSFO with names from over 1000 ontologies. To add cross-references (Xrefs) only to relevant ontologies, we relied on two metrics: *Coverage*, which determined the percentage of LSFs that exist in the target ontology, and *Overlap*, which assessed the percentage of names from the target ontology found within our LSFO. Based on these criteria, we manually narrowed down our selection to 50 ontologies to which we added Xrefs. This enhances interoperability and enables integration of existing ontologies, for example, by importing all child fine-grained concepts under the matched target name from domain-specific ontologies, furthering the development of a more comprehensive and interconnected LSFO.

The team of ontology curators and corpus annotators is a highly professional team with experience in the field. Specifically, six out of the eight members hold a PhD — and the remaining two an MSc — in bioinformatics or computer science. All authors work actively in the field of biomedical data science, half of them having biomedical NLP as their main research focus and having participated in the creation of several biomedical corpora (Pafilis et al., 2013; Luoma et al., 2023; Nastou et al., 2024; Mehryary et al., 2024; Kim et al., 2009; Pyysalo et al., 2012), and two having previously authored papers related to ontology design (Speer et al., 2018; Hoehndorf et al., 2011).

### 2.5 Dictionary-based NER

The JensenLab tagger (tagger hereafter) (https://github.com/larsjuhljensen/tagger) is a fast dictionary-based NER system that recognizes a wide variety of biomedical entities based on underlying dictionaries (Jensen, 2016). To provide a dictionary-based NER system for LSF recognition, we integrated the tagger into our workflow, enhancing its functionality by supplying a dedicated LSF dictionary. The LSF dictionary was constructed using names from LSFO, alongside orthographic variant generation and a block list to enhance recall and precision, respectively. The tagger assigns a single unique identifier to synonyms and automatically generated name variations (e.g. plural and adjective forms) of the same LSF, enabling effective NEN for matched names. To improve precision, we use a block list to exclude problematic names that would cause many false positives during tagging. Specifically, we manually inspected all names that gave more than 2000 matches in 36.1 million PubMed abstracts (as of August 2023) and 4.5 million articles from the PMC open access subset (as of April 2022) to identify those that should be added to the block list.

### 2.6 Transformer-based NER

To explore the potential of using transformers for NER, we adapted an existing NER system (Luoma *et al*., 2023). Specifically, we built upon the RoBERTa-large-PM-M3-Voc model (RoBERTa-bio hereafter), which has demonstrated the best performance in several NER tasks (Lewis *et al*., 2020; Miranda-Escalada *et al*., 2023). We trained the model for multiclass classification of the nine categories of LSFs using LSF200 without OOC annotations. The 200 abstracts were divided into a 40-document holdout test set and a 160-document combined train and development set. We made sure this split was balanced across the nine categories of LSFO. Hyperparameter selection was done through a grid search using a 5-fold cross-validation setup. This was done instead of a simple train–development split to avoid the risk of overfitting on a small development set. To determine the best hyperparameters for the model’s final evaluation, we computed the micro F1-score based on the total True Positives (TP), False Positives (FP), and False Negatives (FN) across all classes and folds. Finally, we trained a model on all available training and development data with the selected optimal hyperparameters and evaluated it on the holdout test set.

## 3 Results and Discussion

### 3.1 The LSF ontology

Being able to identify and categorize concepts as diverse as LSFs was the main challenge of this work and it was achieved through the creation of a comprehensive and inclusive LSF ontology (LSFO). Table 1 presents an overview of LSFO, with the last column displaying the number of Xrefs per LSF category. One of the most important and well-studied categories of LSFs is Nutrition. To get more fine-grained concepts for this category, we incorporated food groups from the widely used FoodOn ontology. Table 1 presents both statistics with and without expansions from FoodOn.

**Table 1.** LSFC statistics. In parentheses, the numbers including expansion with terms from FoodOn are shown for the *Nutrition* category and the total number of entries in LSFC.

| Category | Unique LSFs | Total names | Xrefs |
| --- | --- | --- | --- |
| Nutrition (expanded with FoodOn) | 766<br>(4710) | 987<br>(5126) | 2093<br>(4875) |
| Socioeconomic factors | 302 | 604 | 994 |
| Environmental exposures | 175 | 694 | 643 |
| Substance use | 104 | 283 | 596 |
| Physical activities | 84 | 171 | 283 |
| Personal care products and cosmetic procedures | 78 | 139 | 361 |
| Non-physical leisure time activities | 58 | 124 | 68 |
| Sleep | 45 | 104 | 129 |
| Mental health practices | 40 | 78 | 135 |
| Total (expanded with FoodOn) | 1652<br>(5597) | 3184<br>(7324) | 5302<br>(8097) |

### 3.2 The LSF200 Corpus

LSF200 comprises 200 abstracts, with a total of 39,416 tokens based on BERT basic tokenization (https://github.com/spyysalo/bert-vocab-eval). It contains 1876 manually annotated mentions of LSFs, which are broken down into categories in Table 2. The majority of the LSF mentions fall into the categories *Nutrition, Socioeconomic factors, and Physical activities*. In contrast, the categories *Non-physical leisure time activities, Mental health practices, and Personal care products and cosmetic procedures* are less prevalent in LSF200. The distribution of names is consistent with the statistics of names in the different LSF categories of LSFO (Table 1), showcasing that despite selecting journals to represent all nine categories of LSFs, names from the largest categories tend to appear in most abstracts. Evaluating the quality of the manual annotations in terms of interannotator agreement gave an F1-score of 83%. This score emphasizes the challenging nature of annotating concepts as diverse as LSFs, making it difficult to formulate a comprehensive definition and achieve perfect annotation even at a human level.

**Table 2.** LSF200 corpus statistics.

| Category | Total mentions | Unique names | % of names per category |
| --- | --- | --- | --- |
| Nutrition (expanded with FoodOn) | 445 | 270 | 27.36% |
| Socioeconomic factors | 275 | 174 | 17.63% |
| Environmental exposures | 199 | 132 | 13.37% |
| Substance use | 187 | 75 | 7.60% |
| Physical activities | 222 | 88 | 8.92% |
| Personal care products and cosmetic procedures | 56 | 35 | 3.55% |
| Non-physical leisure time activities | 61 | 27 | 2.74% |
| Sleep | 110 | 57 | 5.78% |
| Mental health practices | 74 | 29 | 2.94% |
| OOC | 247 | 100 | 10.13% |
| Total | 1876 | 987 | 100% |

We also introduced a special category of annotations called Out of context (OOC), which we use to highlight mentions that appear in a context that falls outside the scope of LSFs. For example, the word “tobacco” is normally an LSF but not when it appears in the context of “tobacco company”.

### 3.3 Evaluation of dictionary-based NER

To assess the performance of our dictionary-based NER system, a crucial component involves evaluating the dictionary generated from the LSFO. It is important to note that this dictionary is not constructed using the text corpus, and, as such, it is appropriate to employ the entire LSF200 for evaluation of the dictionary and NER system.

We present performance results in two versions: one with OOC mentions included in the annotated corpus and one without. This evaluation assesses the impact of removing OOC mentions on the performance of dictionary-based NER. In the initial version, OOCs are treated as typical LSF mentions, resulting in an F1-score of 65.2% (precision: 96.0%, recall: 49.4%). In the second version, we removed OOC mentions entirely from annotations. The resulting performance was a slightly lower F1-score of 63.6% (precision: 85.0%, recall: 50.8%). When OOCs are not treated as LSFs, the number of false positives increases considerably, due to the presence of the OOC names in the dictionary, which results in a large drop in precision from 96.0% to 85.0%. Conversely, how OOC mentions are counted has very little impact on recall.

Figure 2 shows the performance for each of the nine categories, revealing that the recall varies much more than the precision and that OOC has a similar impact on precision across categories. For most categories, OOC mentions have only a minor impact on recall, with “Non-physical time activities” being the only notable exception. For this category, many OOC mentions go undetected by the dictionary-based NER system, and the recall thus improves substantially when OOC mentions are excluded.

**Fig. 2.**
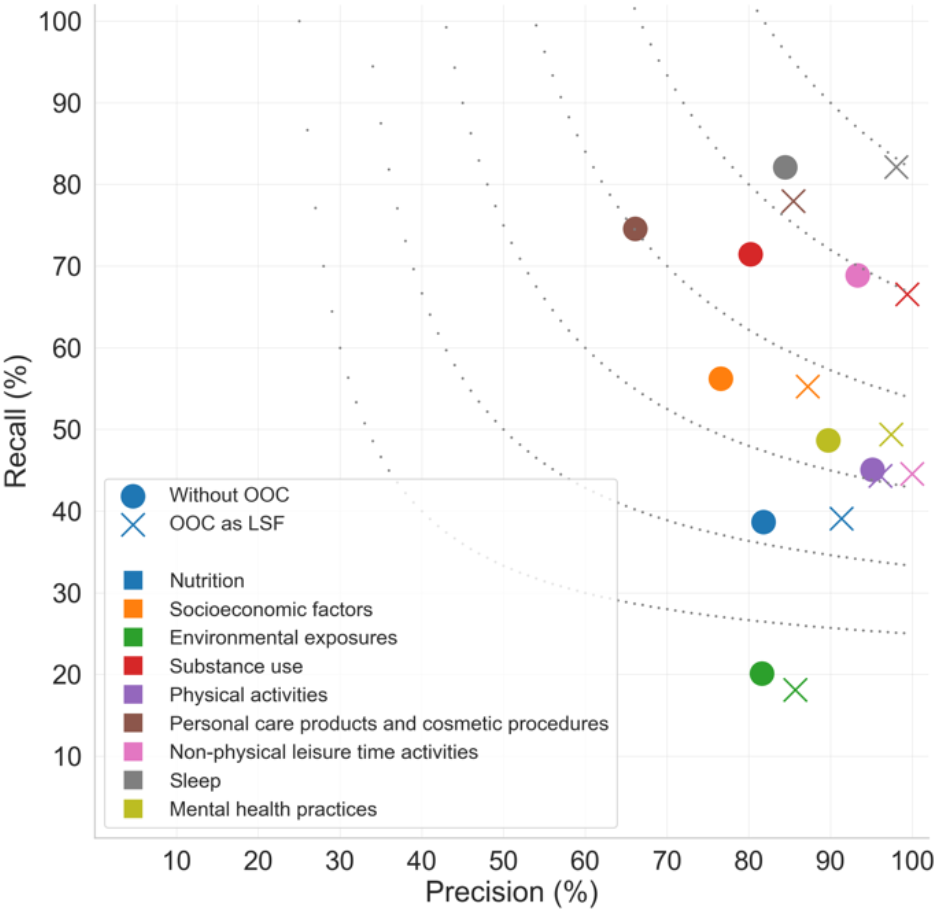
Performance of dictionary-based NER system in two OOC annotation variants per LSF category.

The wide range of recall values is explained by fundamental differences between the categories. For example, the dictionary by design does not have concepts for all possible exposures, for which reason the “Environmental exposures” category has very low recall. This implies that the low recall can likely be improved by using the existing Xrefs to integrate finegrained concepts from, e.g., the Exposome Explorer.

In LSF200, this issue was not immediately evident, as it is selected from journals where all these entities are represented as LSFs. However, in a real-world scenario where the entire literature is tagged, this approach could result in detrimental effects on precision. Detailed performance results for various categories are available in Supplementary Section 6.

#### 3.3.1 Manual error analysis

As anticipated, the dictionary-based NER exhibits impressive precision but lower recall due to its limitations in recognizing out-of-dictionary mentions. In Table 3, we have categorized the missing LSF mentions (false negatives) by the dictionary-based NER system.

**Table 3.** Error analysis for dictionary-based NER (False Negatives)

| Error category | Total mentions | Unique names |
| --- | --- | --- |
| Abbreviation & Synonym | 332 | 155 |
| Ambiguous name | 213 | 97 |
| Fine-grained concept | 211 | 128 |
| Discontinuous entity | 20 | 14 |
| Annotation error | 17 | 15 |
| Total | 793 | 409 |

The majority of missing LSF mentions are due to either abbreviations or synonyms related to existing LSF concepts in the dictionary. This is promising, as it suggests LSFO covers most of the relevant concepts, and that the recall of the NER system can thus be improved by enriching our dictionary with more synonyms for existing concepts. Other false negatives are due to ambiguous names that have been intentionally excluded; for example, “therapy” can denote both psychotherapy, which falls under mental health practices, or medical treatments, which are not considered LSFs. This approach prioritizes precision by avoiding matches with too many irrelevant hits, although it results in missing some relevant mentions in LSF200.

The last major group of missing LSF mentions is “Fine-grained LSF concepts”, which refers to mentions that cannot be classified as major missing LSF concepts because the parent LSFs of these mentions already exist in the dictionary. The “Environmental exposures” is, as already mentioned, a prominent example of this. The fact that missing concepts in LSFO account for only 211 errors in 1876 total LSF mentions, and that all of these are fine-grained concepts, suggests that the ontology has high quality in terms of its breadth.

The final few false negatives observed are either due to discontinuous entities, which cannot be identified by a dictionary-based method (e.g. “Vitamin C” in the expression “Vitamins E and C”) or simply annotation errors made by the human curators and are thus not actual errors of the NER system.

As the dictionary-based system has high precision, it produces much fewer false positives than false negatives, the majority of which are due to OOC mentions as already described (Table 4). The remaining few false positives are either due to dictionary errors (names that should not have been in the dictionary), ambiguous names (which could be blocked at the price of more false negatives), or annotation errors in the corpus.

**Table 4.**
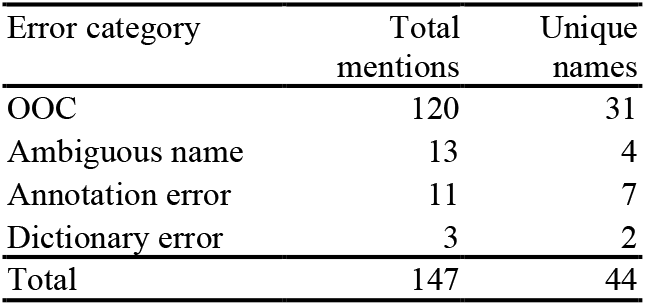
Error analysis for dictionary-based NER (False Positives)

### 3.4 Comparison with transformer-based NER

To compare the performance of dictionary-based NER to state-of-the-art methods, we trained a transformer-based system on LSF200 without OOC mentions (see Methods for details). We evaluated the system on the 40document test set of LSF200. The overall NER performance was 70.1% precision and 85.3% recall, corresponding to an F1-score of 77.0%. To allow direct comparison, we also reevaluated the dictionary-based NER system on the test set only, yielding 76.6% precision, 53.8% recall, and 63.2% F1-score. The results clearly highlight that the dictionary-based NER maximizes the precision by exclusively tagging predefined names from the dictionary and utilizing a block list to avoid tagging problematic names. In contrast, the transformer-based NER achieves better recall due to its ability to detect LSF mentions based on the context. Figure 3 shows the performance of dictionary-based NER and transformer-based NER for the test data within each LSF category.

**Fig. 3.**
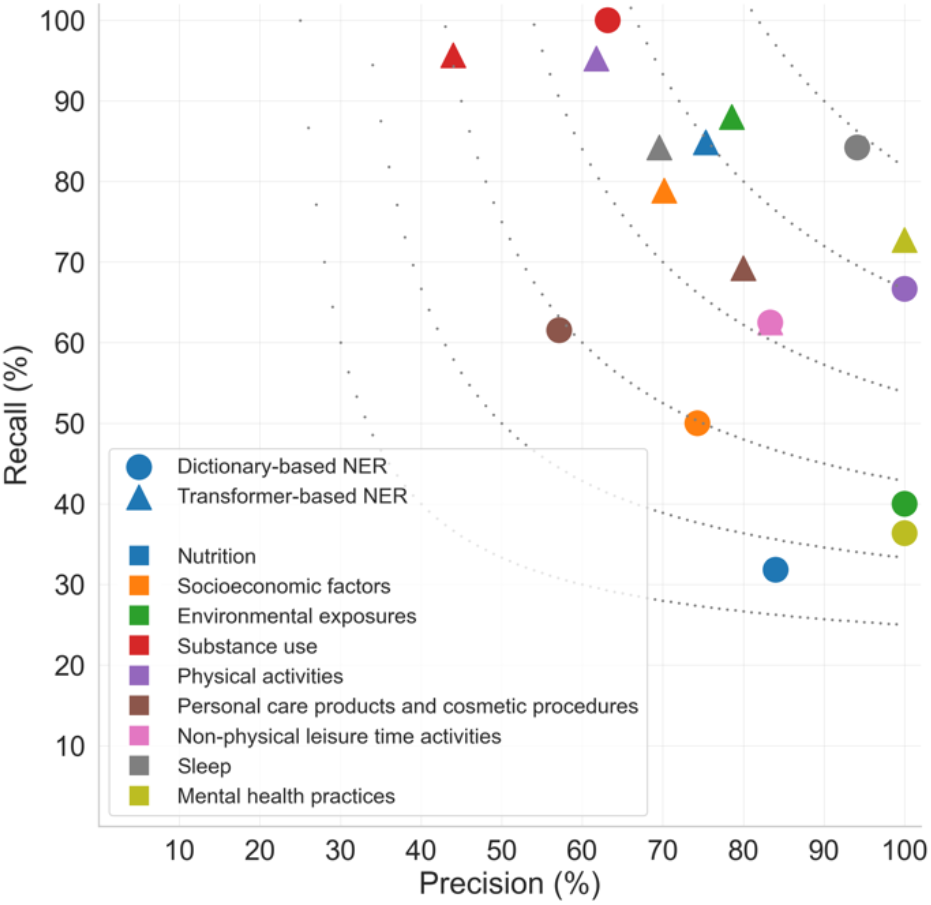
Performance of dictionary-based NER and Transformer-based NER only for test data per LSF category.

The transformer-based NER demonstrates promising recall across most categories, with *Nutrition, Mental health practices*, and *Socioeconomic factors* showing particularly large improvements in recall with only small drops in precision. For *Personal care products and cosmetic procedures*, transformer-based NER surprisingly improved mainly the precision. Finally, the transformer-based NER system performs worse than the dictionary-based one in terms of both precision and recall for categories *Substance use* and *Sleep*.

#### 3.4.1 Manual error analysis

In Table 5 we categorize the errors produced by the Transformer-based NER, encompassing both names missed by the system (FNs) and names incorrectly detected as LSFs (FPs). The system also makes a few mistakes on discontinuous entities, although much fewer than the dictionary-based system.

**Table 5.**
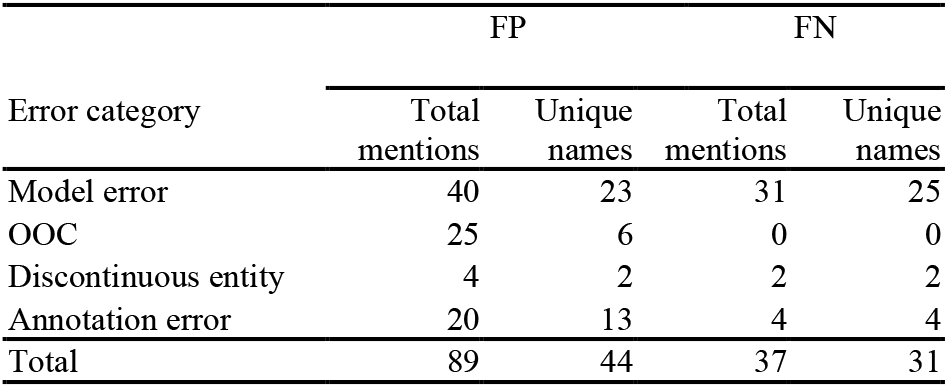
Error analysis for Transformer-based NER.

The majority of the errors, however, were labeled as “Model error”, encompassing the broad class of cases where the model, for no obvious specific reason, fails to provide accurate predictions. Lastly, we have again some errors, which upon inspection turn out to be mistakes in the manual annotation rather than the model being wrong.

### 3.5 Large-scale tagging of the scientific literature

Results for the tagging of 36.1 million PubMed abstracts (as of August 2023) and 4.5 million articles from the PMC open access subset (as of April 2022) using both the dictionary-based NER and the transformer-based NER have been made available via Zenodo (https://zenodo.org/records/10450308). There are in total 85,919,460 LSF matches for dictionary-based NER, with approximately half of them (44%) corresponding to *Nutrition* terms and a quarter (23%) to *Socioeconomic factors*. Tagging with the transformer-based system yielded 251,607,761 total matches. *Nutrition* terms once again constitute almost half of the matches of the system (48%), and *Environmental exposures* come second at 19%. Figure 4 shows the matches per LSF category for the two methods. A key difference between the results of the two systems is that the dictionary-based system inherently provides matches that are normalized to LSFO identifiers whereas the transformer-based system does not. This makes the former a better starting point for many other textmining tasks such as relation extraction.

**Fig. 4.**
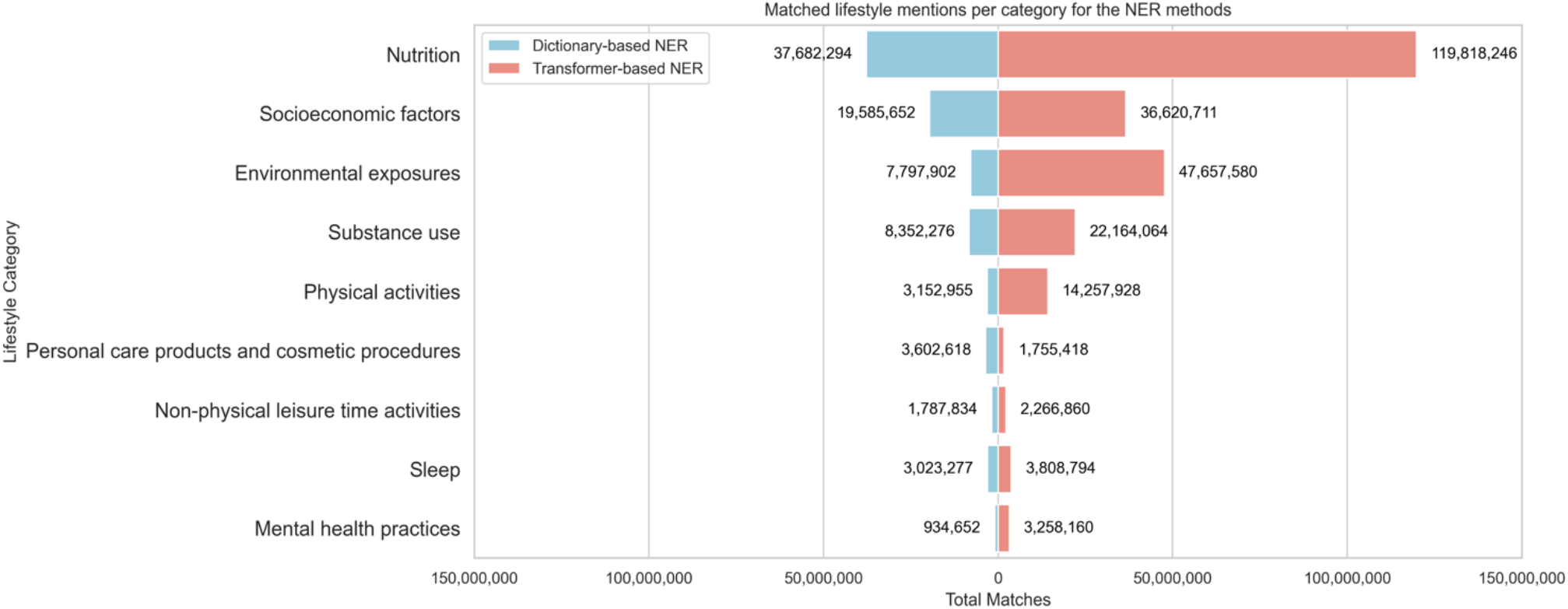
Matched lifestyle mentions from large-scale tagging of scientific literature using the NER methods

## 4 Conclusions

In this paper, we introduce novel resources to address the challenge of recognizing LSFs within biomedical text. We present dictionary-based NER and transformer-based NER systems, both demonstrating promising performance in identifying LSFs. The LSF Classification stands out as a diverse and hierarchical classification of LSFs that we used as a backbone for the dictionary-based NER system, but which can serve as a resource for standardizing LSF information in general. Furthermore, the manually annotated LSF200 corpus proved to be sufficient for training a transformer-based NER and for evaluating both types of NER systems. The presented NER systems, LSFO, LSF200 corpus, and matched LSFs from large-scale runs of both NER systems on PubMed and PMC-OA articles, are made publicly available under open licenses to facilitate further research.

## Supplementary data

Supplementary data are available at Bioinformatics online.

## Funding

This work was supported by the Novo Nordisk Foundation [NNF14CC0001, NFF17OC0027594]. K.N. has received funding from the European Union’s Horizon 2020 research and innovation program under the Marie Sklodowska-Curie [101023676]. M.K. has received funding from Novo Nordisk Foundation LSFC[NNF20SA0035590].

## Conflict of Interest

none declared.

## Notes

https://github.com/EsmaeilNourani/LSFO-expansion

### Competing Interest Statement

The authors have declared no competing interest.

### Summary of Updates

We have renamed the introduced resource name from LSFC to LSFO and added relevant explanations about the provided ontology

