## Supplementary for "Lifestyle factors in the biomedical literature: An ontology and comprehensive resources for named entity recognition"

### 1. LSF200 text corpus

To create an LSF text corpus, we selected three of the most relevant journals within each of the nine LSF categories. We selected 200 abstracts for our corpus, evenly distributed across all categories. We aimed for a balanced selection of documents among LSF categories, considering the chosen journals' scope and focus. The table below shows a list of journals per category which are considered for random document selection.

List of journals per category for the selected 200 documents

| Lifestyle-factor branch | Journal Names | Publisher |
|---|---|---|
| Environmental exposures | Environment International | Elsevier |
| | Indoor Air | Wiley |
| | Journal of the Air & Waste Management Association | Taylor & Francis |
| Nutrition | International Journal of Food Sciences and Nutrition | Springer |
| | European Journal of Nutrition | Springer |
| | Plant Foods for Human Nutrition | Springer |
| Physical activities | Journal of Physical Activity and Health | Human Kinetics Publishers Inc. |
| | Sports Medicine | Springer |
| | Journal of Aging and Physical Activity | Human Kinetics Publishers Inc. |
| Mental health practices | Psychotherapy Research | Taylor & Francis |
| | International Journal of Group Psychotherapy | Taylor & Francis |
| | American Journal of Psychotherapy | American Psychiatric Association |
| Socioeconomic factors | Journal of Interpersonal Violence | Sage Journals |
| | The European Journal of Contraception & Reproductive Health Care | Taylor & Francis |
| | Child Abuse & Neglect | Elsevier |
| Personal care products and cosmetic procedures | Journal of Dental Hygiene | American Dental Hygienists' Association |
| | Journal of Orthodontics | Sage (British Orthodontic Society) |
| | Journal of Orofacial Orthopedics | Springer |

| | | |
|---|---|---|
| Substance use | Nicotine & Tobacco Research | Oxford |
| | Drug and Alcohol Dependence | Elsevier |
| | Tobacco Control | BMJ Journals |
| Non-physical leisure time activities | Journal of Gambling Studies | Springer |
| | Cyberpsychology, Behavior, and Social Networking | Mary Ann Liebert, Inc. |
| | Journal of Physical Activity and Health | Human Kinetics Publishers Inc. |
| Sleep | Sleep Medicine Reviews | Elsevier |
| | Sleep | Oxford |
| | Journal of Sleep Research | Wiley |

### 2. Guidelines for manual creation of the first version of the LSFs dictionary

The initial manual version of the dictionary comprised 9 categories: Nutrition, Socioeconomic Factors, Environmental Exposures, Substance use, Physical Activities, Non-physical leisure time activities, Beauty and Cleaning, Sleep, and Mental Health Practices. Below you will find the initial guidelines that were used to manually add terms in the first version of the dictionary. It should be noted that the dictionary has undergone significant restructuring from this original version to generate the ontology currently available online: https://github.com/EsmaeilNourani/Lifestyle-factors-classification/tree/db7df9e0ab5137430079ee1e14bc04684be6bd34

#### Nutrition

Encompasses some general terms related to nutrition, e.g. "malnutrition", and five main sub-categories, with more sub-categories: Dietary habits, Food groups, Food processing and preparation methods, Macronutrients and Micronutrients. The sub-categories Food groups, Macronutrients and Micronutrients were inspired from nutritools.org.

- **Dietary habits** include **"dieting schemes**", e.g. **"low fat"** diet or **"Asian diet"**; the child term "**dietary restriction**" refers to dieting schemes that impose the exclusion of one or more food groups from a person's diet, e.g. "**vegan diet**". **"Meal planning"**, **"meal frequency"** and various **"eating behaviors"**, e.g. **"emotional eating"**, are also included in the dietary habits.
- **Food groups:** Includes foods commonly stated in questionnaires. The hierarchical organization was inspired by FoodOn (https://foodon.org/) and FooDB (https://foodb.ca/).
- The sub-categories of **Macronutrients** and **Micronutrients** were inspired by https://mynutrition.wsu.edu/nutrition-basics and https://www.nal.usda.gov/fnic .
- **Food processing and preparation:** The child term **"cooking methods"** refers to household or restaurant cooking, whereas the term **"thermal processing of food"** refers to industrial food processing.
- **Food packaging**: Includes types and subtypes of packaging

#### Socioeconomic factors

Preliminaries:

- Definitions (from Wikipedia):
    - **income:** For households and individuals income is the sum of all the wages, salaries, profits, interest payments, rents, and other forms of earnings received in a given period of time – also known as gross income. Net income is defined as the gross income minus taxes and other deductions (e.g., mandatory pension contributions).
    - **wealth:** Wealth is the abundance of valuable financial assets or physical possessions which can be converted into a form that can be used for transactions. [...] Wealth refers to the value of everything a person or family owns. This includes tangible items such as jewelry, housing, cars, and other

personal property. Financial assets such as stocks and bonds, which can be traded for cash, also contribute to wealth.

- ❏ **socioeconomic status:** Socioeconomic status (SES) is an economic and sociological combined total measure of a person's work experience and of an individual's or family's economic and social position in relation to others. When analyzing a family's SES, the household income, earners' education, and occupation are examined, as well as combined income, whereas for an individual's SES only their own attributes are assessed. [...] When placing a family or individual into one of these categories, any or all of the three variables (income, education, and occupation) can be assessed.
- ❏ **social status**: "Social status" is a measurement of social value. More specifically, it refers to the relative level of respect, honor, assumed competence, and deference accorded to people, groups, and organizations in a society. Some writers have also referred to a socially valued role or category a person occupies as a "status" (e.g., gender, social class, ethnicity, having a criminal conviction, having a mental illness, etc.)."

- Social stratification and the distinction of **"social class"** and **"social status"** was based on the three-component theory of stratification, (a.k.a. **Weberian stratification** or the three class system): According to the German sociologist Max Weber, **class**, **status** and **power** are distinct ideal types. Weber developed a multidimensional approach to social stratification that reflects the interplay among wealth, prestige and power. Weber argued that power can take a variety of forms. A person's power can be shown in the social order through their status, in the economic order through their class, and in the political order through their party:
    - **Wealth (Class)**: includes property such as buildings, lands, farms, houses, factories and as well as other assets – Economic Situation
    - **Prestige (Status)**: the respect with which a person or status position is regarded by others – Status Situation (e.g. *poets and saints can possess immense influence on society with often little economic worth*)
    - **Power**: the ability of people or groups to achieve their goals despite opposition from others – Parties (e.g. *an employee of the Federal Bureau of Investigation, or a member of the United States Congress, may hold little property or status, but they still hold immense power*)

Socioeconomic factors in the dictionary include:

- **"personal relationships"**: Refers to any type of relationship a person can have with another person (romantic, friendly and family relationships). Family relationships also include the way someone was raised, e.g. **"divorced parents",** "**parent-child conflict".**
- **"residing location":** Captures the difference between living and/or growing up in a **developed** *vs.* **developing country**, **urban** *vs.* **rural area** *etc.*
- **"religion"**: Whether someone is religious or/and was raised in a religious family. **"Religious upbringing"** is also a child of **"family relations and upbringing"** under the **"personal relationships"** sub-category.
- **"education"**: The classification of the **"education level"** sub-category was done according to the International Standard Classification of Education (ISCED) of 2011.

- **“occupation”:** Includes types of occupation, such as **“blue collar workers”, “white collar workers”, “sedentary work”** that may also be interconnected.
- **“Social class”** division into **“underclass”, “lower class”, “middle class”** and **“upper class”** was done based on the “Three-level economic class model” .

Socioeconomic status is typically broken into three levels (high, middle, and low) to describe the three places a family or an individual may fall into. When placing a family or individual into one of these categories, any or all of the three variables (income, education, and occupation) can be assessed.

#### Environmental exposures

The child term **“exposures related to the place of residence”** includes temporary exposures, e.g. **“forest fire”**, as well as more permanent “**environmental conditions”,** such as **“climate and weather conditions”. “Food chain contamination”** may be related to the place of residence or not, thus it is both under **“exposures related to the place of residence”** and **“Environmental exposures”**. Environmental exposures also include the large sub-category of **“xenobiotics”** (in short chemicals that accumulate in organisms due to contamination). The terms included in “xenobiotics” are a result of brainstorming and google search, cross-validated and enriched by Exposome Explorer (http://exposome-explorer.iarc.fr/classifications/5287).

#### Substance use

Refers to use of illicit/recreational drugs, not those prescribed by doctors for medical conditions, as well as terms related to “**smoking**”. It also includes general terms that refer to substance use, e.g. **“recreational drugs”**, **“drug usage”**. Good online sources for drug classification include:

- https://www.greenfacts.org/en/psychoactive-drugs/index.htm#1
- https://www.verywellmind.com/what-is-psychoactive-22500
- https://www.addictioncenter.com/drugs/drug-classifications/central-nervous-system-depressants/
- https://www.drugbank.ca/categories/DBCAT000437

#### Physical activity

Includes physical activity related specifics, such as **“exercise frequency”, “exercise intensity”, “exercise type”. “Exercise type”** includes the **“aerobic exercise”** and **“anaerobic exercise”** sub-categories with examples of relevant activities for both, e.g. **“jogging”** and **“crossfit”.** Aerobic and anaerobic exercise are essentially related to the exercise intensity, as the production of ATP in the mitochondria by the use of oxygen is sufficient to meet the energy demands.

Links between exercise type and exercise intensity:

  - https://www.medicinenet.com/aerobic_exercise/article.htm
  - https://www.emedicinehealth.com/aerobic_exercise/article_em.htm
  - https://en.wikipedia.org/wiki/Aerobic_exercise

Apart from the physical activity related specifics, physical activity is divided into 4 main categories: physical activity performed in the leisure time, physical activity required in a person's occupation, e.g. "**construction workers**", physical activity required in the household, e.g. **"vacuum cleaning"**, as well as means of transportation that require physical activity, e.g. **"cycling".** The **"leisure time physical activity"** is divided into **"group physical activities"**, i.e. activities normally done in teams, such as **"playing football",** and **"individual physical activities",** such as **"running".** Of course the "individual physical activities" could be also performed in groups, but that is neither a requirement nor the norm, as it is with the examples under the "**group physical activities**".

#### Non-physical Leisure Time Activities

Includes any activity an individual might be preoccupied with in their free time that is not a physical activity, e.g. music, books, but also traveling or going out. The sub-category **"use of electronic devices"** encompasses all activities that require the use of an electronic device with a screen, such as **"video games"**, **"watching TV"** and **"mobile phone use"**. The latter term has the sub-category **"smartphone use",** which in turn has the sub-category **"social media use"** that finally collapses to **"facebook"**, **"instagram"** and **"social media addiction"** in general. The terms included are the combined result of brainstorming and inspiration from https://www.questionpro.com/survey-templates/leisure-time-activities/

#### Beauty and Cleaning

Refers to "**hygiene"** habits (e.g. **"shower daily"**), "**use of cosmetic and cleaning products"**, as well as invasive procedures that people undergo to improve their appearance, such as **"cosmetic surgery**". **"Cleaning products"**, such as **"bleach"** or **"oven cleaners"**, are commonly considered toxic to humans. **"Personal care products"** may also contain chemicals that are (controversially) associated with human diseases, e.g. the association of **"aluminum chlorohydrate"** or **"parabens"** with breast cancer (https://www.cancer.org/cancer/cancer-causes/antiperspirants-and-breast-cancer-risk.html). Note that the list was the result of brain-storming, therefore it is not exhaustive.

#### Sleep

Based on brainstorming, PubMed research and the Stanford Healthcare sleep questionnaire.

#### Mental health practices

Mental health practices refer to the activities, strategies, and approaches used to maintain, improve, and support an individual's mental and emotional well-being. The practices included herein are a result of brainstorming and the following sources:
https://www.nimh.nih.gov/health/topics/caring-for-your-mental-health
https://www.nhs.uk/mental-health/talking-therapies-medicine-treatments/talking-therapies-and-counselling/types-of-talking-therapies/
https://medlineplus.gov/howtoimprovementalhealth.html

### 3. Dictionary expansion from Wikipedia titles

The overall workflow is presented in Supplementary Figure1 and explained step by step below.

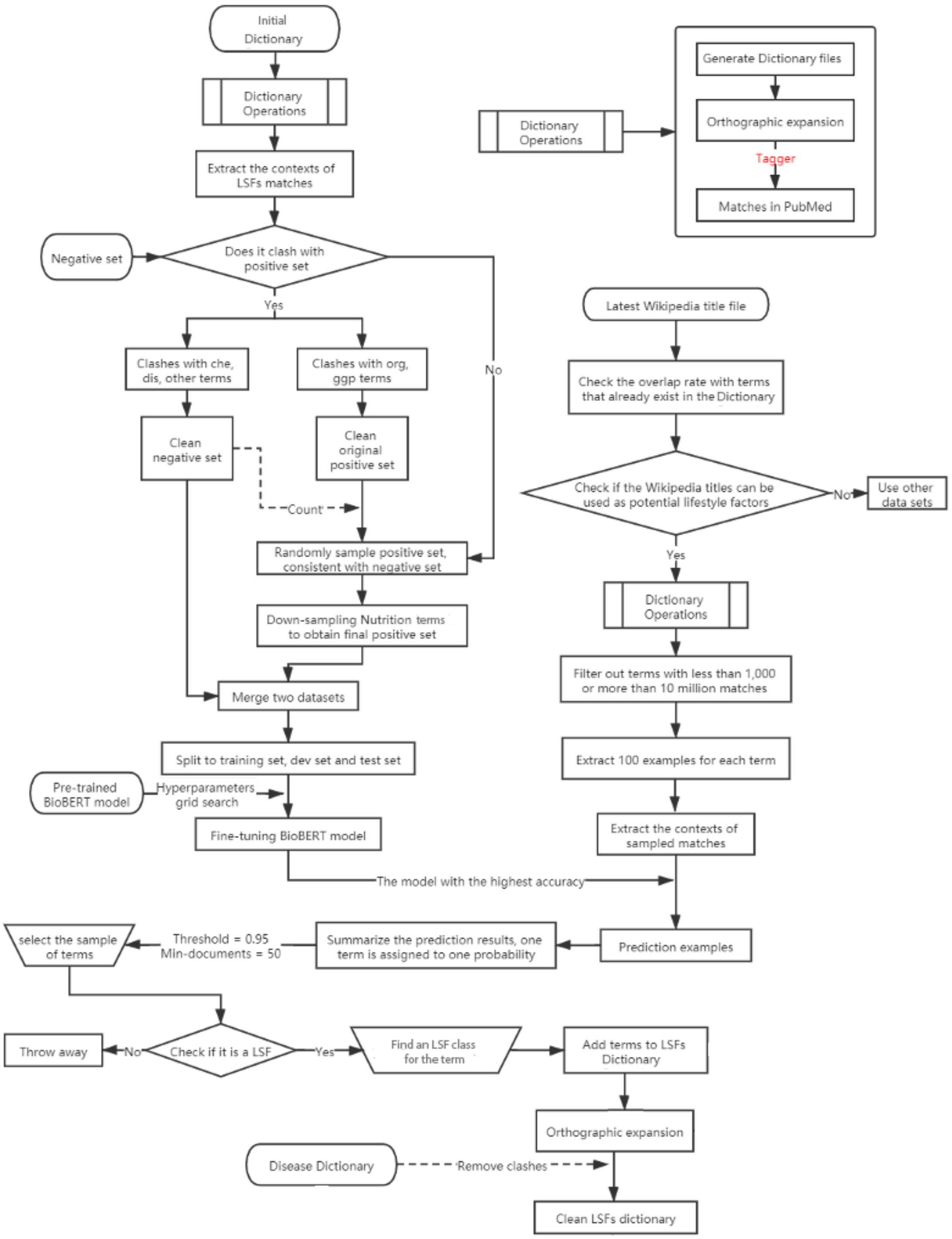


Supp. Fig.1: The overall workflow for LSF dictionary expansion from Wikipedia titles

Starting from the top left, the initial manually constructed dictionary described in Supplementary Section 1, the biomedical literature, questionnaires, and brainstorming was used to generate dictionary files, do orthographic expansion (Pafilis *et al.*, 2013), and run the

Jensenlab tagger (Jensen, 2016) against the entire biomedical literature to get all matches for LSFs. This was then used to extract contexts around these matches and create an initial positive class (positive corresponds to LSF) to train the BioBERT-based entity type classifier introduced by Nastou *et al.*, 2023. In order to train the classifier the same number of negative examples needed to be introduced in the training set. As an initial dataset from which examples were extracted the small dataset introduced in this work (Nastou *et al.*, 2023) was used, which contained contexts around five entity types, namely genes, diseases, species, chemicals, as well as, negative nouns and noun phrases (other) for these four classes. To ensure that there are no clashes with the positive LSF set, the negative set was meticulously cleaned as shown in the workflow. This step ensures that the negative set comprises only terms that are unambiguously negative with respect to the concept of interest.

Once the negative set was free of conflicts the number of total examples (count) was used to generate the positive set, with an 1:1 ratio of positive to negative. Extra caution was taken so that the positive set is balanced in terms of how many terms in the positive examples come from the different classes of LSFs. This step is crucial, particularly for nutrition terms, to prevent any class imbalance that could later bias the Transformer-based model training. Finally, the positive set was merged with the negative set to form a comprehensive dataset of ~486,000 examples in total, which is available via Zenodo (https://zenodo.org/records/10450308). This dataset was further divided into three subsets: the training set (70%), the development set (20%), and the test set (10%). These subsets were used to fine-tune a pre-trained BioBERT model. Hyperparameter optimization was performed through grid search to identify the model configuration that yields the highest accuracy. The table below shows the sets of parameters used during grid search. The best model reached an accuracy of 94.61% and is available through Zenodo (https://zenodo.org/records/10450308). Its hyperparameters were: maximum sequence length=256, batch size=32, learning rate=3e-5, number of epochs=2. This model was used in the subsequent prediction phase.

| Model | $BioBERT_{Base}$ |
|---|---|
| Maximum Sequence length | 96, 128, 256 |
| Batch Size | 16, 32, 64 |
| Learning Rate | 5e-5, 3e-5, 2e-5, 5e-6 |
| Number of Epochs | 1,2 |

The right part of the workflow shows how Wikipedia titles were used to augment the dictionary. The procedure begins by checking for term overlap between Wikipedia titles and existing terms in the initial dictionary. The aim was to identify if that data source is relevant, which was the case as 45% of the terms in the initial dictionary are Wikipedia titles (1827/4019).

The workflow proceeded with generating a dictionary from Wikipedia titles to tag the entire PubMed using the Jensenlab tagger. The Wikipedia title dictionary and the tagger matches results are available through Zenodo (https://zenodo.org/records/10450308). Titles that were too rare (with less than 1,000 matches) or too common to be actual LSFs (more than 10 million matches) were filtered out to maintain a focus on relevant and manageable datasets, as the final set of names would need to be manually checked before addition to the dictionary. 100 examples were extracted for each of the remaining titles to facilitate the predictive phase, where these were classified as lifestyle factors (positive) or not (negative). The probabilistic scores produced by the classifier for each example were aggregated and a final score for each title was generated using the same process as Nastou *et al.*, 2023.

At this time, since the screening process was manual, a probability threshold was required as the criterion for inclusion for Wikipedia titles. In order to select an appropriate threshold, the terms that already exist in the LSFs ontology needed to be excluded from the results file. After that, 20 titles were randomly selected at 6 threshold intervals, as shown in the table below. These were manually inspected and the frequency of potential LSFs among them was calculated. When the threshold was set to >0.95, 35% of the terms were manually classified as LSFs, and after that, the frequency of LSFs dropped. For this reason, an aggregated score threshold of 0.95 and a minimum number of documents where the term appears of 50 were chosen for compiling a list of names to be potentially included in the dictionary after manual inspection.

| Score threshold interval | Frequency of LSFs |
|---|---|
| >0.98 | 0.25 |
| >0.95 | 0.35 |
| 0.90~0.95 | 0.2 |
| 0.85~0.90 | 0.15 |
| 0.8~0.85 | 0.15 |
| 0.75~0.80 | 0.1 |

448 out of the 2144 terms that surpassed the score threshold were manually classified as LSFs and added to the LSF dictionary. The LSF dictionary was again subjected to orthographic expansion to include all relevant variations of the newly added terms. Then the dictionary underwent a cleaning process where terms that clash with diseases were removed. This ensures that the two dictionaries remain distinct and accurate, without any overlapping terms that could interfere with their future potential use in biomedical text mining. Finally, the dictionary was further manually cleaned to ensure its robustness, geared towards enhancing the extraction of lifestyle factors from the biomedical literature.

### 4. Dictionary expansion from diverse knowledge bases and ontologies

In the subsequent phase of dictionary/ontology expansion, LSF-related candidates were extracted from several resources.

Statistics of LSF candidates used for dictionary/ontology expansion.

| Resource | Number of LSF Candidates | |
|---|---|---|
| | Extracted | Unique |
| BioPortal | 139322 | 56843 |
| Wikidata | 55119 | 40203 |
| DBPedia | 48861 | 29794 |
| ConceptNet | 30260 | 13797 |
| WordNet | 5688 | 2681 |
| Total | 279250 | 143318 |

LSF candidates are assigned a probability score based on a combination of two distinct scoring methods.

The first score measures the degree of semantic similarity between the candidate and existing LSFs in the dictionary, thereby emphasizing the identification of synonyms. To accomplish this, we convert the existing LSF names into corresponding 768-dimensional dense vectors using a sentence transformer (https://huggingface.co/sentence-transformers/all-mpnet-base-v2), to accurately capture the semantics of phrases that are not single words. Then, we create an efficient and scalable index (https://github.com/spotify/annoy) using the vectors of all existing LSF names to search for the nearest neighbors within this index for each candidate. For candidates that have very close neighbors in this index, it is probable that they are not new LSF concepts but synonyms of existing ones. Additionally, we created an index using non-LSF names to serve as a negative sample set. Each candidate was evaluated for its proximity to terms in both the LSF and non-LSF indices. We calculated scores based on the inverse square of the mean distance to the nearest neighbors in each index. These scores were then normalized to probabilities, indicating the likelihood of a candidate being an LSF.
The second score covers broader scope and not only synonyms using a topic modeling

approach called BERTopic (Egger and Yu, 2022), which can identify concepts related to LSF topics. BERTopic is a topic modeling technique that leverages pre-trained transformer-based language models to create topics. In this study, BERTopic is trained in a semi-supervised fashion where both labeled existing LSFs, non-LSF names and unlabeled candidates are considered for training the topic model. The advantage of using unlabeled candidates for training the model is that we can identify new lifestyle-related topics which do not exist in the current ontology and thereby identify new LSF concepts and not only synonyms. To train the topic model, we incorporate the text surrounding each sample. This method allows us to capture both the embeddings derived from sample names and the contextual variations associated with the samples. The topic model identifies existing topics within the training set, encompassing both LSF and non-LSF topics. Subsequently, we manually select LSF topics to score candidates using the trained model. For each candidate, the likelihood of being an LSF is calculated as the cumulative sum of probabilities assigned by the model, indicating the candidate's association with all previously shortlisted LSF-related topics.

Finally, to score candidates using the combined scores from the two aforementioned approaches, we calibrate them to ensure the score values are comparable. The calibrated scores are then used to calculate a consensus for scoring and filtering the candidates. We selected the top-scoring candidates and added them to the ontology after manually checking and potentially filtering out irrelevant ones.

The code is available at https://github.com/EsmaeilNourani/LSFO-expansion.

### 5. Ontology construction and conflict resolution

In the ontology construction process, as explained in Supplementary Section 1, clear guidelines were established for the different categories of lifestyle factors (LSFs) and what should be considered an LSF for each category. These guidelines made it easier for ontology creators to evaluate each candidate and determine whether it qualified as an LSF and which branch it belonged to.

We preserved the hierarchy of concepts from FoodOn to include all fine-grained concepts under the "Food Groups" branch. For concepts queried from other ontologies, integration was done selectively after manual curation, without necessarily preserving the hierarchy of the source ontology. The extracted and scored LSF-related candidates were presented to the curators for them to determine if they should be added to the ontology and, if so, where they should be placed.

To streamline the curation process and ensure curators had all the necessary information to make informed decisions, we developed a dashboard that provides the following information for each candidate LSF:

- A definition of the candidate LSF from Wikipedia, if available, along with a sample image
- The option to automatically do a Google search for the candidate LSF
- Similar existing LSFs in the ontology, based on semantic similarity as explained in Supplementary Section 3
- Automatically queried sample sentences from PubMed, showing how the candidate LSF appears in the literature, with the option to search for additional examples
- The local structure of the ontology for the selected target LSF, making it easier to decide on the correct placement for the new concept

This tool facilitated ontology creation by providing all the necessary information for making accurate decisions and included a mechanism to collect feedback and comments from curators. If the two primary curators agreed on where a candidate LSF should be added in the ontology, the action was performed automatically. If they disagreed, two additional experienced curators decided whether to add it or not. These cases were discussed in regular meetings involving the main curators and the two experienced curators. The latter also performed a final review of the ontology.

### 6. Evaluation of dictionary-based NER

Performance of dictionary-based NER system in two OOC annotation variants per LSF category.

| | Without OOC | | | OOC as LSF | | |
|---|---|---|---|---|---|---|
| Lifestyle-factor branch | Precision | Recall | F | Precision | Recal | F |
| Environmental exposures | 81.63 | 20.1 | 32.26 | 85.71 | 18.1 | 29.89 |
| Nutrition | 81.82 | 38.65 | 52.5 | 91.36 | 39.07 | 54.73 |
| Physical activities | 95.15 | 45.05 | 61.14 | 96.12 | 44.3 | 60.65 |
| Mental health practices | 89.74 | 48.65 | 63.09 | 97.44 | 49.37 | 65.53 |
| Socioeconomic factors | 77.13 | 56.36 | 65.13 | 87.23 | 55.24 | 67.64 |
| Personal care products and cosmetic procedures | 67.74 | 75.0 | 71.19 | 85.48 | 77.94 | 81.54 |
| Substance use | 85.63 | 71.66 | 78.02 | 99.4 | 66.53 | 79.71 |
| Non-physical leisure time activities | 93.33 | 68.85 | 79.25 | 100.0 | 44.55 | 61.64 |
| Sleep | 88.35 | 82.73 | 85.45 | 98.06 | 82.11 | 89.38 |
